# Gated Coupling of Dopamine and Neuropeptide Signaling Underlies Perceptual Processing of Appetitive Odors in *Drosophila*

**DOI:** 10.1101/159897

**Authors:** Yuhan Pu, Melissa Megan Masserant Palombo, Yiwen Zhang, Ping Shen

## Abstract

How olfactory stimuli are perceived as meaningful cues for specific appetitive drives remains poorly understood. Here we show that despite their enormous diversity, *Drosophila* larvae can discriminatively respond to food-related odor stimuli based on both qualitative and quantitative properties. Perceptual processing of food scents takes place in a neural circuit comprising four dopamine (DA) neurons and a neuropeptide F (NPF) neuron per brain hemisphere. Furthermore, these DA neurons integrate and compress inputs from second-order olfactory neurons into one-dimensional DA signals, while the downstream NPF neuron assigns appetitive significance to limited DA outputs via a D1-type DA receptor (DoplR1)-mediated gating mechanism. Finally, Dop1R, along with a Gβ13F/Irk2-mediated inhibitory and a Gαs-mediated excitatory pathway, underlie a binary precision tuning apparatus that restricts the excitatory response of NPF neurons to DA inputs that fall within an optimum range. Our findings provide fresh molecular and cellular insights into cognitive processing of olfactory cues.

## Summary

Olfaction is an ancient sense that is vital to the survival of animals across evolution. For example, to successfully search for distant or hidden energy sources in an ecological niche, a forager must be able to selectively recognize diverse food-related volatiles among others and evaluate their potential reward values. In addition, olfaction is also known to be functionally interconnected to other senses such as taste, and it may significantly impact cognitive functions that facilitate food detection and competition^1–4^. Therefore, elucidation of conserved neural substrates and mechanisms underlying cognitive processing of food odor stimuli in behaving animals is likely to provide valuable insights into the inner workings of the brain and its evolution.

Numerous studies have revealed that insects and mammals appear to share remarkable anatomical and functional similarities at multiple levels, including odor detection by olfactory receptor neurons, wiring schemes for relaying sensory inputs to second-order projection neurons as well as the presence of analogous spatial maps encoding processed odor information in the higher-order processing center (e.g., the lateral horn in flies and the cortical amygdala in mice that controls innate olfactory behaviors) ^5,6^. However, despite of such extensive knowledge about the sensory processes within the initial two steps of the olfactory circuit, how olfactory information is represented to deeper brain regions and translated to motivated behaviors remains largely unclear.

Genetically tractable *Drosophila* larvae have an advanced yet numerically simpler nervous system. We recently developed a fly larva model that is potentially useful for studying higher-order processing of food-related odor stimuli^7^. In response to a brief presentation of a banana-like scent or balsamic vinegar vapor, well-nourished fly larvae have been shown to display an aroused appetitive state in anticipation of sugar food. In this work, we show that in response to food-related odors *Drosophila* larvae display macronutrient-specific appetitive arousal in an inverted U manner. By taking genetic, pharmacological and neurobiological approaches, we show that perceptual recognition of appetitive odor stimuli requires a neural circuit comprising an ensemble of four DA neurons and a downstream neuropeptide F (NPF, a sole fly homolog of mammalian neuropeptide Y or NPY) neuron per brain hemisphere. These DA neurons, previously shown to form synaptic connections with second-order olfactory (projection) neurons ^7^, integrate and compress olfactory inputs from projection neurons to one-dimensional DA outputs. The NPF neuron functionally interacts with the upstream DA neurons by selectively assigning appetitive significance to a narrow spectrum of incoming DA signals via a D1-type DA receptor (Dop1R1)-mediated gating mechanism. We also provide evidence that Dop1R, along with a Gβ13F/Irk2-mediated inhibitory and a Gαs-mediated excitatory pathway, define a binary precision tuning apparatus in the NPF neuron that restricts the host cell response to DA inputs within an optimum range. Together, our findings suggest that the fly larva is a useful animal model for mechanistic studies of perceptual and other cognitive processing of olfactory and possible other sensory cues.

## Results

### Attribution of anticipated food reward to discrete odor stimuli

Well-nourished *Drosophila* larvae display a persistent state of appetitive arousal in anticipation of sugar food after a brief stimulation of food odors ^7^. Such odor-aroused food anticipation in individual larvae can be demonstrated, following the removal of the odor stimulus, by measuring the contraction rate of external mouth hooks that larvae use to scoop glucose-containing liquid food into the oral cavity ^7,8^ To investigate how food-related olfactory stimuli of varied chemical property and strength are perceptually processed, we first evaluated larval appetitive responses to three chemotactically attractive monomolecular odorants, pentyl acetate (PA), heptanal (Hep) or trans-2-hexenal (T2H) at various doses^9^. We found that while the effective dose range of each odorant differed significantly, their appetizing effects invariably followed an inverted-U function (Figure 1A-C). In addition, balsamic vinegar vapor, a chemically complex odor mixture, also exhibited inverted-U effects (Figure 1D).

**Figure 1.**
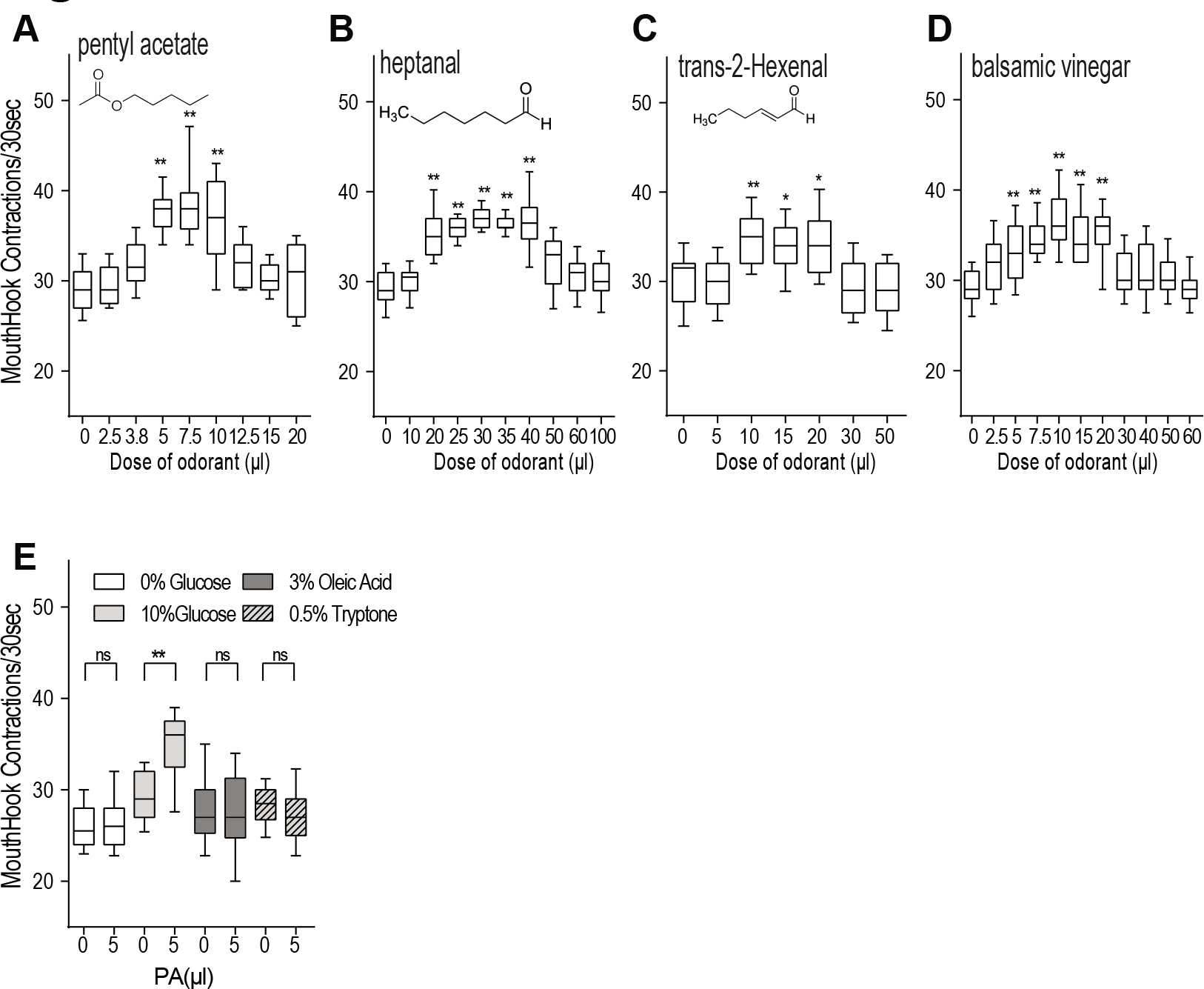
Attribution of anticipated food rewards to discrete odor stimuli by fed larvae. **(A-D)** Third-instar fed larvae (74h after egg laying, AEL) were exposed to an odor stimulus for 5 minutes in a sealed chamber, fumigated with defined concentrations of monomolecular odorants PA, Hep, T2H or natural odorants balsamic vinegar, prior to the feeding test in 10% glucose liquid media. The odor effects on larval feeding rate were quantified by counting mouth hook contractions of each larva over a 30s period. The rate of mouth hook contraction is positively correlated to the amount of dyed food ingested ^7^ n≥15; **(E)** Feeding responses of fed larvae to various types of food after stimulation by an effective dose of PA (5μl). n≥8; Sidak’s multiple comparisons test was used. All behavioral quantifications for this and other figures were performed under blind conditions. Unless indicated otherwise, all the statistical analyses were performed using One-way ANOVA followed by a Dunnett’s test. *P<0.05; **P<0.001. The following figure supplement is available for figure1: Figure supplement 1. Control larvae display normal feeding responses after heat shock treatment

We also examined whether fed larvae, aroused by a particular appetitive stimulus (e.g., 5μl PA), display motivational specificity to a selected type(s) of macronutrients. In addition to 10% glucose medium, 3% oleic acid (unsaturated fatty acid) and 0.5% tryptone (digested protein) diets are also palatable to fly larvae ^10,11^, as evidenced by similar baseline feeding activities of fed larvae in each food medium. However, when stimulated by PA, fed larvae exhibited an increased feeding response to glucose but not oleic acid and tryptone media (Figure 1E). Therefore, these results suggest that fly larvae are endowed with an innate ability to selectively extract salient features from diverse food-related odor stimuli based on their chemical properties and strengths and to differentially attribute a specific type(s) of anticipated food reward to a particular olfactory cue.

### DA neuronal responses to appetizing and non-appetizing odor stimuli

How do fly larvae perceptually judge whether an odor stimulus is appetizing or not? To address this question, we turned our attention to a cluster of DA neurons (named DL2-1 to 4 in each brain lobe) that are essential for larval appetitive response to PA ^7^; Figure 2A). By taking advantage of the fluorescent Ca^2+^ sensor named CaMPARI (Calcium Modulated Photoactivatable Ratiometric Integrator) ^12^, we quantitatively analyzed the excitatory responses of the DL2 neurons to an effective dose of an odorant(s) in behaving fed larvae. The odor stimulations were performed under the same conditions as described for feeding behavioral assays, followed immediately by 5-sec irradiation with 405 nm blue light. This light irradiation irreversibly turns Ca^2+^-bound CaMPARI protein from green to red fluorescence, thereby capturing the excitatory state of the DL2 neurons in the aroused larvae. We observed that stimulation by an appetizing dose of PA (7μl) induced excitatory responses from the four DA neurons in each cluster, as evidenced by increased red fluorescence signals in the DL2 neurons over a 5-min test period (Figure 2B). Similarly, an appetizing dose of balsamic vinegar (20 μl) also triggered a similar rise of intracellular Ca^2+^ levels in the DL2 neurons during its stimulation.

**Figure 2.**
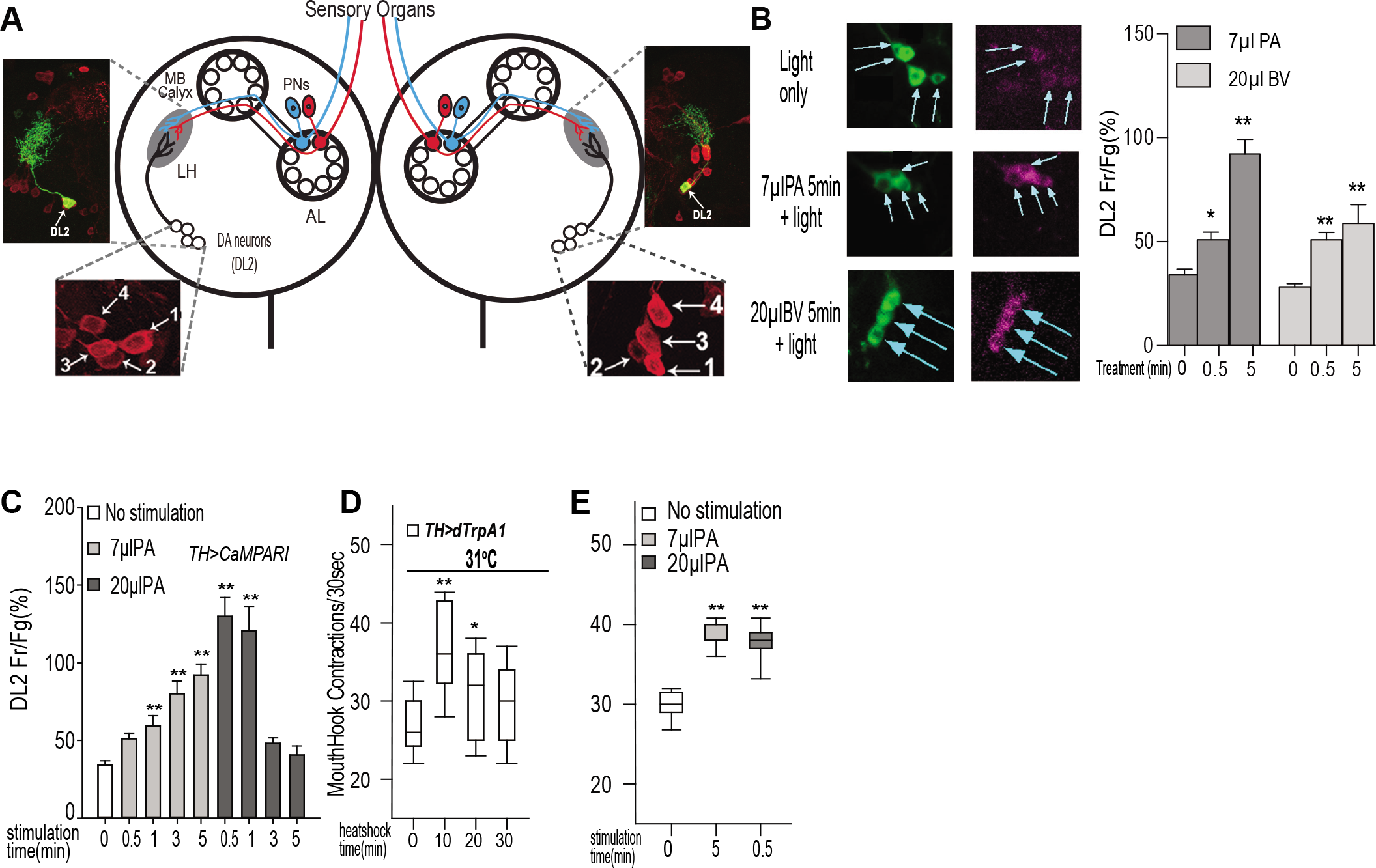
Differential responses of four DA neurons to appetizing and non-appetizing odor stimuli. **(A)** Schematic drawing of the larval olfactory circuit shows two clusters of four DA neurons (DL2-1 to 4) in the left and right brain hemisphere^7^. The DL2 neurons project dendrites to the lateral horn (LH) in each brain lobe, where they form synaptic connections with the projection neurons. Upper insets show that the axons and dendrites of individual DL2 neurons, labeled with mCD8::GFP, are exclusively located in the LH region. AL: antenna lobe; MB: mushroom bodies; PNs: projection neurons. **(B)** CaMPARI-based fluorescence imaging of freely behaving larvae revealed the dynamic responses of four DL2 neurons to an effective dose of PA or balsamic vinegar (BV) during 5-minute stimulation. For each time point, at least 29 DA neurons were imaged. **(C)** In freely behaving larvae, DL2 neurons display distinct dynamic responses to an appetitive dose (7μl) or non-appetitive dose (20μl) of PA, as revealed by CaMPARI-based fluorescence imaging. Data are presented as ±SEM. n≥7; **(D)** Genetic activation of fed larvae expressing the dTrpAl transgene at 31°C for 10 minutes led to increased feeding response in the absence of odor stimulation (n≥19). Also see Figure supplement 1. **(E)** Fed larvae display appetitive responses to 30-sec stimulation by 20μl PA, which is ineffective when stimulation time is 5 min (see Figure 1A). n≥11; *P<0.05; **P<0.001

In parallel, we also examined how the DL2 neurons respond to an ineffective dose of PA in fed larvae under the same experimental conditions. In response to 20μl PA stimulation, DL2 neurons showed a rapid surge of intracellular Ca^2+^ within the first minute, followed by a gradual reduction of the Ca^2+^ increase as the stimulation time is prolonged (Figure 2C). Therefore, the ineffectiveness of 20μl PA to arouse appetitive behavior may be accounted for by two different explanations. One possibility is that 20μl PA may cause the silencing of DL2 neurons during the 5-minute stimulation, thereby disrupting acutely required DA signaling to downstream targets. Alternatively, its stimulation may trigger an excessive release of DA, which is functionally ineffective. To distinguish the two possibilities, we genetically activated the DL2 neurons over different time periods using dTrpA1, a temperature-sensitive TRP family cation channel^13^. A brief activation of these neurons triggered an increased larval feeding response in the absence of odor stimuli, while the prolonged activation still failed to do so (Figure 2D; also see Figure supplement 1). In addition, fed larvae stimulated by 20μl PA for 30 seconds instead showed a significant increase in their feeding response (Figure 2E). Together, these results support the notion that an excessive release of odor-evoked DA signals rather than silencing of the DA neurons is responsible for the ineffectiveness of higher doses of odors.

### Quantitative influences of endogenous DA levels on the appetizing effects of odor stimuli

Since an excessive release of odor-evoked DA likely underlies the ineffectiveness of stronger odor stimuli, we wondered whether reduction of the endogenous level of DA in fed larvae could increase the dosage required for appetitive arousal. To test this hypothesis, we performed RNA interference (RNAi)-mediated knockdown of the activity of Tyrosine Hydroxylase (TH), a rate-limiting enzyme for DA synthesis ^14^. The fly *ple* gene encodes Tyrosine Hydroxylase. Indeed, *TH-GAL4*/UAS-*ple^RNAi^* fed larvae showed significantly increased appetitive responses to higher doses of odorants (e.g., 20μl PA) that are normally ineffective, but they failed to respond to normally effective doses of odorants (e.g., 5μl PA; Figure 3A). In a separate pharmacological experiment, we pre-fed normal larvae with food containing a TH inhibitor, 3IY, for 4 and 6 hours. The 3IY-treated larvae failed to show appetitive response to a normally effective dose of PA (Figure 3B). In parallel, we also tried to increase the endogenous DA level by pre-feeding larvae with L-Dopa, a dopamine precursor, for 2 and 4 hours. We found that 4 hour pre-feeding with L-Dopa blocked the appetitive response to a normally effective dose (e.g., 5μl or 10μl PA). Conversely, the same L-Dopa treatment rendered a lower ineffective dose of PA to elicit an appetitive response (Figure 3C).

**Figure 3.**
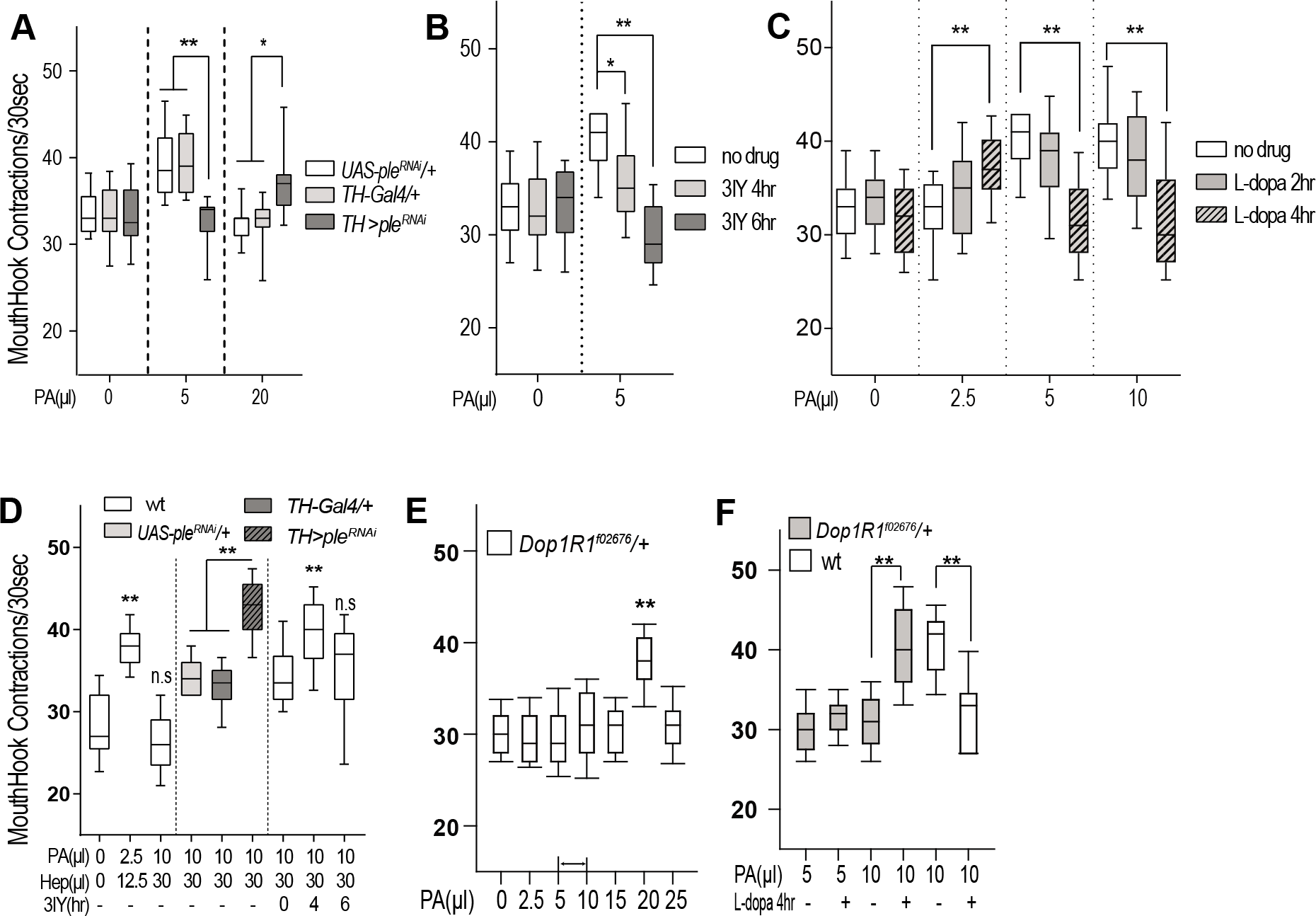
Evidence for Dop1R1-gated attribution of appetitive salience to odor-evoked DA signals. **(A)** Reduced baseline level of DA by expressing UAS-*ple^RNAi^* in TH-GAL4 neurons led to an increase in the minimal effective dose of odorants required for appetitive arousal (n >13). **(B)** Wild type larvae fed with 3IY-containing media for 4 hours blocked the appetitive response to an effective dose of PA (n >11). **(C)** Wild type larvae fed with L-dopa-containing media for 4 hours before feeding tests caused a decrease in the effective dose of odorants required for appetitive arousal (n >10). **(D)** Two odorants in a binary mixture show additive effects. Fed larvae with reduced TH activity showed appetitive responses to a binary mixture containing PA and Hep at effective doses, which is normally not appetitive (n≥10). **(E)** Fed larvae heterozygous for Dop1R1^f02676^, a loss-of-function mutation in Dop1R1, showed a right shift in the inverted-U dose response. Arrows indicate the normal dose range (n≥10). **(F)** Feeding heterozygous Dop1R1^f02676^ larvae with L-dopa restored the appetitive response to a normally effective dose of PA (n≥13). *P<0.05; **P<0.001 The following figure supplement is available for figure3: Figure supplement 2. Heterozygous Dop1R1^f02676^ fed larvae showed right shifts in odor-aroused feeding

Since pre-feeding of L-dopa can augment the appetizing effect of PA at a low dose, we tested whether another lower ineffective dose of odorant (e.g., 12.5μl Hep; see Figure 1B) can be used to substitute the L-dopa treatment for appetitive arousal. Indeed, simultaneous stimulation by 2.5μl PA and 12.5μl Hep was able to significantly increase larval appetitive response (Figure 3D). In contrast, the combined use of two effective doses of PA and Hep was ineffective in appetitive arousal. However, such a binary mixture became effective if the testing larvae have reduced TH-activity or pre-fed with 3IY (Figure 3D). Together, these findings suggest that the minimal strength of an odor stimulus required to arouse appetite is inversely correlated with the baseline level of DA in fed larvae.

### Odor-evoked DA signals acquire motivational salience through a Dop1R1 gating mechanism

Our findings so far have demonstrated the importance of an optimum level of odor-evoked DA signals for larval appetitive arousal. Since the DA receptor Dop1R1 has been shown to mediate the appetitive arousal of fed larvae ^7^, we decided to examine whether Dop1R1 plays a role in presetting the effective dosage of odor-evoked DA signals in fed larvae. Heterozygous *Dop1R1*^f02676^ larvae have a 50% reduction of *Dop1R1* transcripts ^15^. We found that in response to various doses of monomolecular or mixed odorants, *Dop1R1^f02676^*/+ fed larvae displayed a right shift in the dose-response curve (Figures 3E and Figure3--Figure supplement 2). Furthermore, pre-feeding of L-dopa restored their appetitive responses to a normally effective dose of PA (10μl, Figures 3F). Therefore, these findings suggest that Dop1R1 is a major genetic factor that quantitatively delimits the optimum range of effective DA levels, and individual animals with varied Dop1R1 levels likely require distinct levels of odor-evoked DA signals for appetitive arousal.

### Roles of two dorsomedial DA neurons in odor-aroused appetite

Our previous study showed that the *Drosophila* neuropeptide Y-like NPF system contributes to the odor-aroused feeding response of fed larvae by modulating the excitatory responses of the DL2 neurons to olfactory stimuli^7^. Anatomical analyses have revealed that a *npf*-GAL4 driver predominantly labels six NPF neurons in the larval central nervous system (CNS; Figure 4A, B; ^16,17^ Another *npf*-LexA driver, which predominantly labels two dorsomedial NPF neurons in the larval brain, shows that their dendrites are enriched in the lateral horn region (Figure4--Figure supplement 3). To determine which subset(s) of NPF neurons is necessary to induce appetitive arousal, we performed targeted laser lesioning of NPF neurons. We found that odor-stimulated fed larvae missing the two-lesioned dorsomedial NPF neurons failed to display appetitive response, while those larvae with two lesioned dorsolateral NPF neurons showed normal appetitive behavior (Figure 4C).

**Figure 4.**
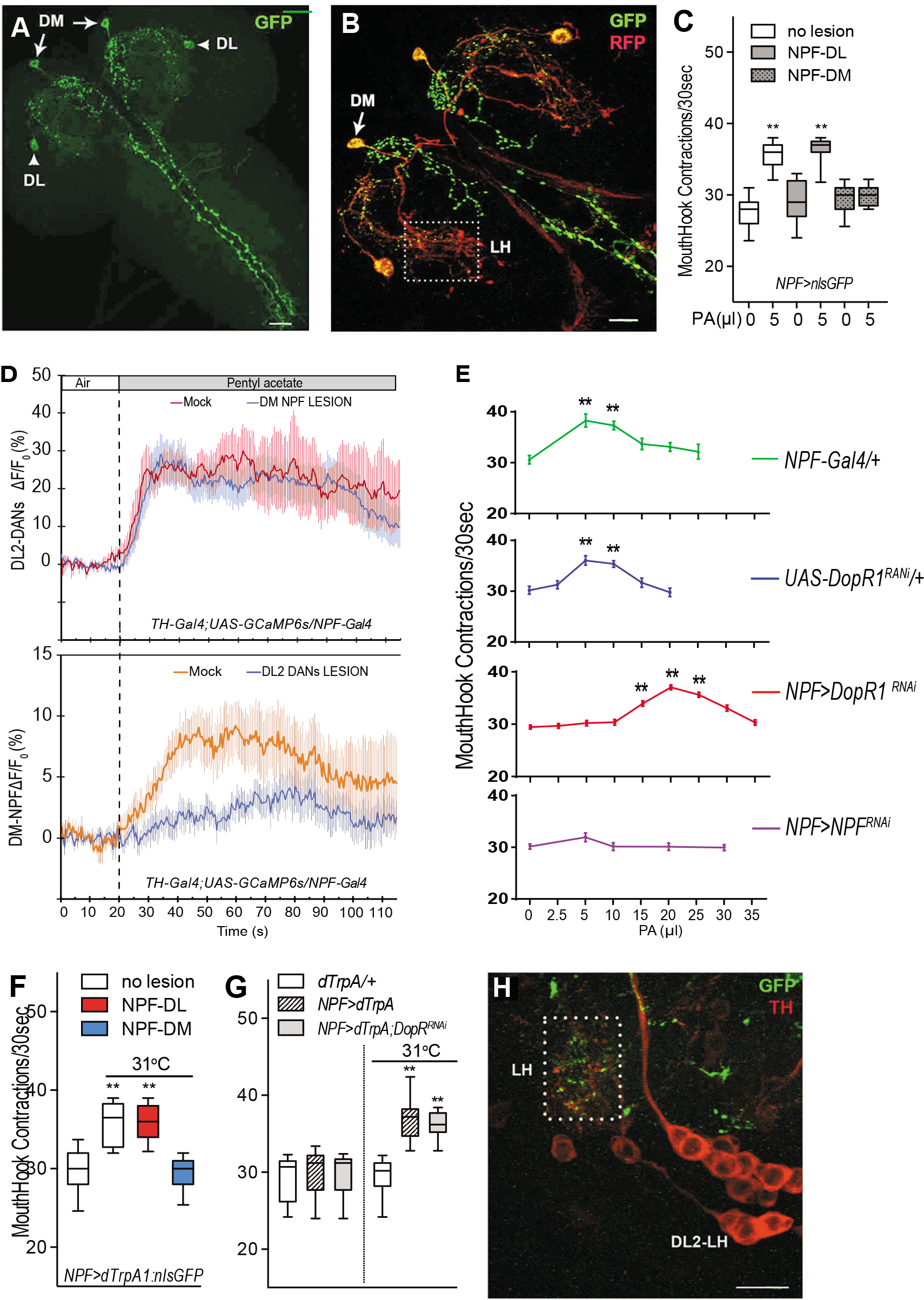
Anatomical and Functional Analyses of Two dorsomedial NPF neurons for Appetitive Arousal. **(A)** Four NPF neurons and the presumptive axons revealed by expression of UAS-nsybGFP^19^ directed by NPF-GAL4. DM: two dorsomedial NPF neurons; DL: two dorsolateral NPF neurons. Two NPF neurons in the subesophageal ganglia are out of the focal plane ^16^. **(B)** The presumptive axons (green) and dendrites (red) of NPF neurons are labeled by sytGFP and Denmark ^43^. The dotted box shows the dendrites from the dorsomedial NPF neuron at the lateral horn (LH). **(C)** Targeted laser lesioning of two dorsomedial NPF neurons (NPF-DM) but not dorsolateral NPF neurons (NPF-DL) in NPF-GAL4/UAS-nlsGFP fed larvae abolished the appetitive response (n>10). **(D)** Functional influences of targeted lesioning of the NPF or DA neurons on their responses to PA stimulation over a 90-sec period. **(E)** Functional knockdown of Dop1R1 activity in NPF neurons led to a right shift in the dose-response curve, while the *npf* knockdown abolished larval appetitive responses to PA at all doses tested (n>15). **(F)** Genetic activation of NPF neurons in NPF-GAL4/ UAS-*dTrpA1* fed larvae at 31°C for 15min led to increased appetitive response (n≥17). Also see related data in Figure4--Figure supplement 3. NPF-GAL4/ UAS-nlsGFP/UAS-dTrpA1 fed larvae with lesions in two dorsolateral NPF neurons displayed normal appetitive response after the heat shock, while the same larvae with two lesioned dorsomedial NPF neurons failed to display appetitive response (n>10). **(G)** Fed larvae expressing dTrpA1 in normal or Dop1R1-deficient NPF neurons showed similar feeding responses (n≥16). **(H)** Presumptive synaptic connections between NPF and DL2 neurons at the LH (the dotted box) are revealed using a split GFP technique^19^, involving *TH-GAL4, NPF-LexA, UAS-CD4::spGFP^1-10^ and LexAop-CD4::spGFP^11^*. DL2-LH: four DL2 neurons projecting to the LH. Green: immuno-fluorescence of split GFP; Red: anti-TH. Scale bar=20pm. *P<0.05; **P<0.001. The following figure supplements are available for figure4: Figure supplement 3. The odor-aroused feeding responses of Dop1R1- or NPF-deficient fed larvae **Video 1.** A 3D view of the presumptive synaptic connections between DA and NPF neurons.

Given that NPF signaling was previously implicated in modulation of DL2 neuronal responses to the outputs from second-order projection neurons, we decided to determine the observed activity of the dorsomedial NPF neuron could be explained by its positive regulation of the upstream DA neurons. To test this possibility, we used a fluorescent Ca^2+^ indicator, GCaMP6 to analyze the excitatory responses of four DL2 neurons to PA stimulation over a 60-sec period in the presence or absence of two dorsomedial NPF neurons. We found that in both cases, the odor responses of DL2 neurons were largely similar. We also performed a reciprocal imaging test under the same odor treatment. In the presence of the DL2 neurons, the NPF neurons were excited by the PA stimulation, but their odor responses were significantly attenuated when the two clusters of DL2 neurons were lesioned (Figure 4D). We also observed that the DL2 neurons responded to the PA stimulation within a few seconds. However, the response of the NPF neurons showed a significant time lag of up to 25 seconds. These results raise the possibility that the dorsomedial NPF neurons may contribute to a yet uncharacterized circuit mechanism by acting downstream from odor-responsive DL2 neurons.

To test this hypothesis, we functionally knocked down Dop1R1 activity in *npf*-GAL4/*Dop1R1^RNAi^* fed larvae. Similar to *Dop1R1*/+ fed larvae mutant larvae, these *Dop1R1*-deficient larvae displayed a right shift of the dose-response curve, phenocopying (Figures 4E, also see Figure4--Figure supplement 3). Also, fed larvae expressing UAS-*npf^RNAi^* under the direction of *npf*-GAL4 showed no significant appetitive response to odorants (e.g., PA and Hep) at any of the doses tested, revealing an essential role of NPF in arousing larval appetite. Furthermore, we genetically activated NPF neurons using a fly TRP family channel, *dTrpA1* ^18^. The *npf*-GAL4/UAS-*dTrpA1* fed larvae, heat shocked at 31°C for 15 or 30 min, showed increased feeding activity even in the absence of appetitive odor stimuli (Figures. 4F and Figure4--Figure supplement 3). In addition, the *npf*-GAL4/UAS-*dTrpA1* larvae with lesions in two dorsomedial but not dorsolateral NPF neurons failed to show increased feeding response, demonstrating that activation of the two dorsomedial NPF neurons is sufficient to induce appetitive arousal in fed larvae (Figure. 4G). The expression of dTrpA1 in *Dop1R1*-deficient NPF neurons also led to bypassing the requirement of Dop1R1 activity to trigger appetitive arousal in an odor-independent manner (Figure 4H), suggesting that Dop1R1 is involved in modulation of DA-mediated excitation of NPF neurons rather than NPF release. Finally, we performed an imaging analysis using a split GFP technique ^19^. This analysis has revealed that the two dorsomedial NPF neurons likely form synaptic connections with the ipsilateral cluster of DL2 neurons in the lateral horn (Figures 4I; also see Video 1). Together, our findings suggest that the dorsomedial NPF neuron and four upstream DA neurons define a neural circuit for perceptual processing of information encoded by a common pool of odor-evoked DA from the DL2 neurons.

### NPF neuronal responses to appetizing and non-appetizing odor stimuli

NPF has been implicated in encoding motivational states for a variety of goal-directed behaviors in larval and adult flies ^20–24^ We postulate that the dorsomedial NPF neuron may contribute to a food odor-aroused motivational state in fed larvae by selectively assigning appetitive significance to DA signals that fall within a limited range. To test this hypothesis, we examined how the dorsomedial NPF neurons respond to appetizing and non-appetizing odor stimuli. We first examined the potential impact of PA stimulation on the membrane potentials of the dorsomedial NPF neurons in a larval preparation using a fluorescent indicator of membrane potential (Arclight) ^25^. The dorsomedial NPF neurons showed a gradual increase in excitatory response during a 10-minute continuous exposure to the vapor of PA at an effective dose. In contrast, when a higher ineffective dose of PA was applied, no excitatory responses were observed, except for a transient depolarization immediately following the odor application (Figure 5A and B). The responses of the dorsomedial NPF neurons to appetizing and non-appetizing odor stimuli were also analyzed in behaving larvae using CaMPARI-based imaging (Figure 5C). Similar to the Arclight imaging results, these neurons showed an excitatory response to an appetizing dose of PA (7μl), but failed to respond to non-appetizing PA doses that are either too high or too low (e.g., 20 or 3.5μl). Therefore, these results indicate that the inverted-U effects of odor stimuli can also be observed at the level of NPF neuronal excitation. On the other hand, in the Dop1R1-defieinct NPF neurons, the effective dose of PA required to activate the dorsomedial NPF neurons is upshifted from 7μl to 20μl PA.

**Figure 5.**
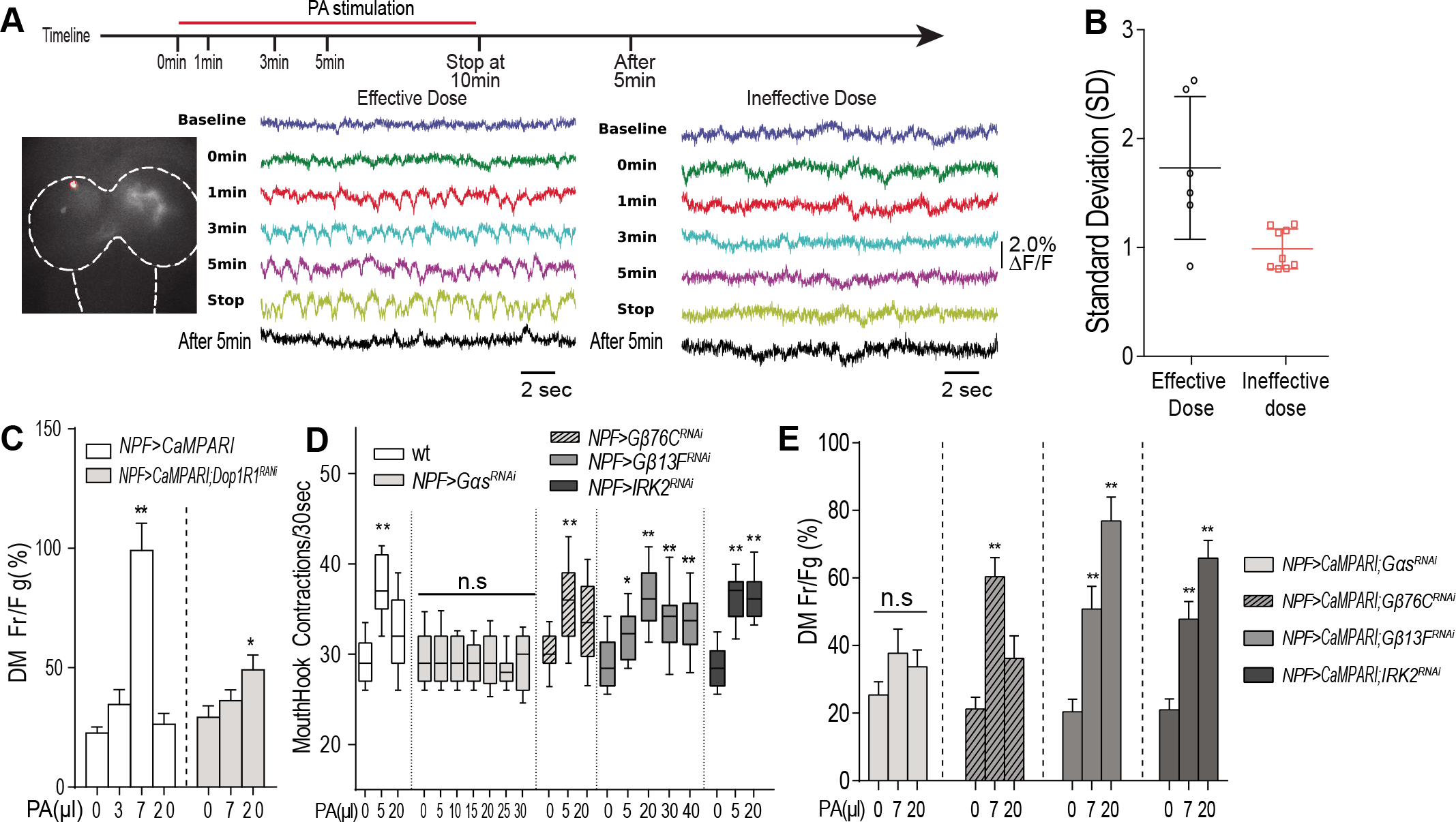
Dop1R1-gated NPF neuronal excitation and an underlying genetic mechanism. **(A)** ArcLight-based imaging of single dorsomedial NPF neurons (circled) in a preparation of NPF-GAL4/ UAS-*ArcLight* fed larvae. Two sample recordings of normalized membrane activities of dorsomedial NPF neurons in two different preparations: one stimulated by an effective dose of PA and the other stimulated by a higher ineffective dose. (B) Standard deviations (SDs) for membrane activity were calculated for both effective (n=6) and ineffective (n=9) treatments at 5 minutes after PA stimulation. The effective odor treatment induced significantly stronger excitatory activities of NPF neurons (t-test, P<0.01). **(C)** CaMPARI-based imaging analysis of dorsomedial NPF neurons in behaving fed larvae. The normal NPF neuron shows an inverted U excitatory response to increasing doses of PA, while the Dop1R1-deficient NPF neurons shows a right shift (n≥6). **(D)** Genetic analyses of the effects of various doses of PA on the appetitive responses of *NPF-Gal4/UAS-Gαs^RNAi^*, *NPF-GAL4/UAS-Gβ13F^RNAi^*, *NPF-GAL4/UAS-Gβ76C^RNAi^*, and *NPF-GAL4/ UAS-IRK2^RNAi^* fed larvae. **(E)** CaMPARI-based imaging analysis of the effects of appetitive and non-appetitive doses of PA on the excitatory responses of Gαs, Gβ76C, Gβ13F or IRK2-deficient NPF neurons (n≥6 for each group). *P<0.05; **P<0.001. The following figure supplement is available for figure5: Figure supplement 4. The effects of functional knockdown of Dop1R1 and its effectors on Odor-aroused Feeding Responses

### A binary genetic mechanism for Dop1R1-mediated precision tuning of NPF signaling

A key remaining question is how Dop1R1 precisely tunes the two dorsomedial NPF neurons to assign appetitive salience to otherwise behaviorally meaningless DA signals. D1-type DA receptor is associated with the heterotrimeric G protein complex consisting of Gαs, Gβ, Gγ subunits ^26^. Upon its activation by DA, the dissociated Gαs subunit and Gβγ complex each defines a separate effector pathway. We found that RNAi-mediated knockdown of Gαs activity blocked appetitive response to PA at all doses tested (5 to 20 μl; Figures 5D, Figure5--Figure supplement 4). Furthermore, in freely behaving fed larvae, the reduced activity of Gαs also significantly attenuated the excitatory response of dorsomedial NPF neurons to an effective dose of PA (Figure 5E).

The *Drosophila* genome contains three separate genes each encoding a different Gβ subunit ^27,28^;. Functional knockdown of one of them, *Gβ13F*, in the NPF neurons of *npf*-GAL4/UAS-*Gβ13F^RNAi^* fed larvae led to an expansion of appetitive response to PA at all doses tested (from 5 to 40μl; Figure 5D, Figure5--Figure supplement 4). Since a G protein-dependent inward-rectify potassium channel could be a potential molecular target of the Gβγ complex ^29^, we functionally knocked down each of the three known IRK genes ^30,31^. Indeed, fed larvae that are deficient in IRK2 but not IRK1 or IRK3 activity phenocopied *npf*-GAL4/UAS-*Gβ13F^RNAi^* fed larvae (Figure 5D, Figure5--Figure supplement 4). In parallel, we also tested how the two dorsomedial NPF neurons in behaving *npf*-GAL4/UAS-*Gβ13F^RNAi^* and *npf*-GAL4/UAS-*IRK2^RNAi^* fed larvae respond to stimulation by 7 or 20 μl PA. We found that the dorsomedial NPF neurons expressing UAS-CaMPARI, along with UAS-*Gβ13F^RNAi^* or UAS-*IRK2^RNAi^*, displayed an abnormally high level of excitatory responses to the stimulation by a normally ineffective dose of PA (20μl), while their responses to 7μl PA remained similar (Figure 5E). These results show that when the Gβ13F/IRK2 pathway is deficient, the effective range of odor-induced DA signals is greatly expanded, converting the inverted U dose-response curve to the sigmoidal shape in these larvae. Together, our findings have revealed a binary genetic mechanism in the NPF neurons that underlies Dop1R1-gated coupling of odor-evoked DA and NPF signaling. Furthermore, Dop1R1 gating of the NPF neurons appears to employ a two-layered precision tuning strategy that ultimately shapes the inverted-U dose response of fed larvae: the Gαs-mediated excitatory pathway is used to set up a minimal threshold level of odor-evoked DA signals required for NPF neuronal excitation; the Gβ13F/IRK2-mediated inhibitory pathway is specialized for preventing NPF neuronal response to any odor-evoked DA signals that are excessively strong (Figure 6).

**Figure 6.**
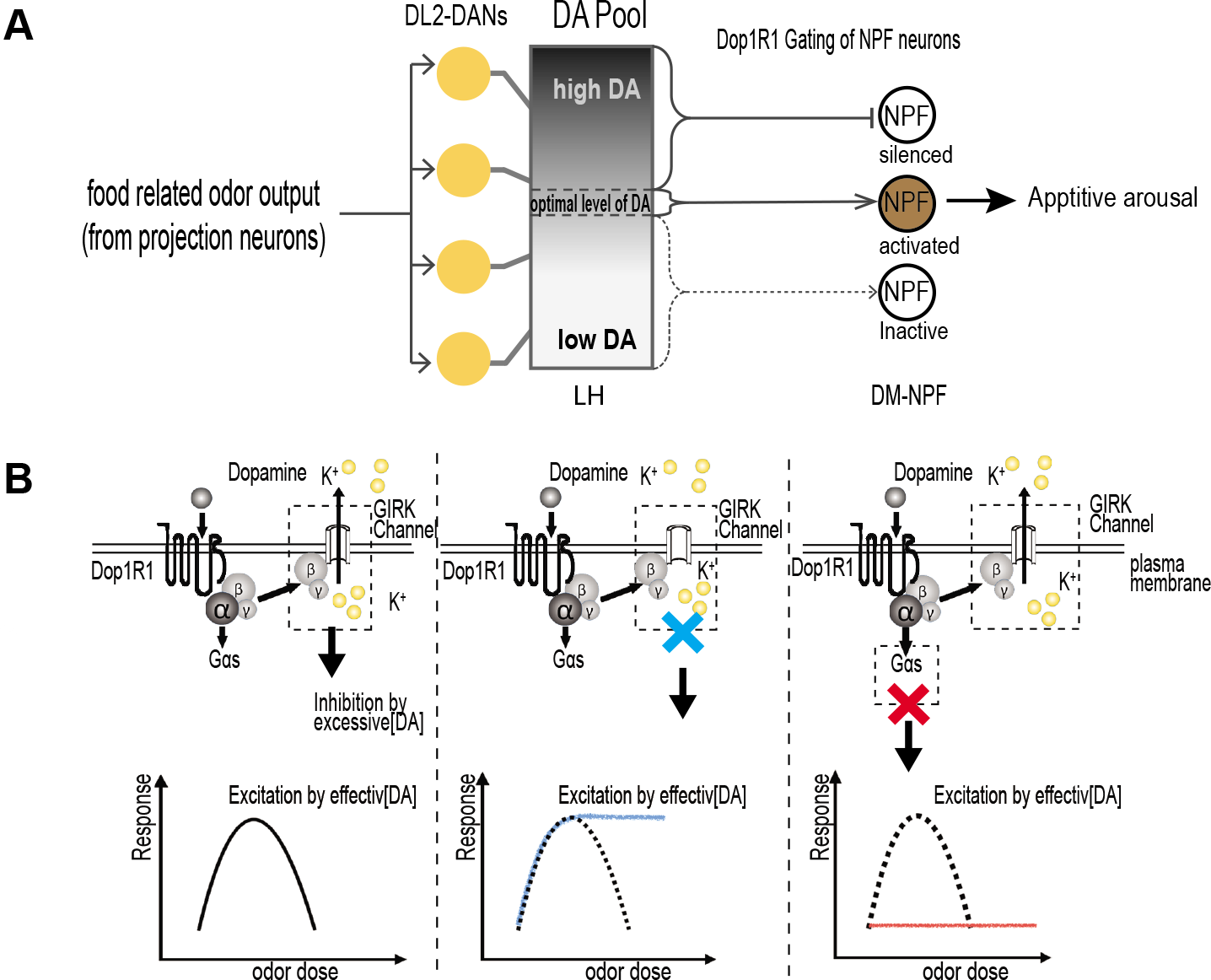
A Working Model for a Neural Network Comprising DA and NPF Neurons and Its Regulation of Appetitive Arousal in Fed Larvae. **(A)** The diagram depicts a perceptual processing module comprising four DA neurons and a postsynaptic NPF neuron in each brain hemisphere of fly larvae. The food-related olfactory outputs from projection neurons are integrated in the DA neurons and transformed to DA signals. A Dop1R1-mediated gating mechanism in the NPF neuron precisely tunes its excitatory response to the optimum levels of DA from a presynaptic pool, resulting in the selective attribution of appetitive significance to a small fraction of odor-evoked DA signals. **(B)** Schematic presentation of a binary genetic mechanism consisting of a Gβ13F/IRK2 inhibitory pathway and a Gαs excitatory pathway. Together, they mediate Dop1R1-dependent precision tuning of the NPF neuron, resulting in its inverted-U excitatory response to odor-evoked DA inputs.

## DISCUSSION

To date, how sensory inputs are perceptually processed in deep brain centers to arouse specific behavioral drives remains poorly understood ^32^. We have shown that *Drosophila* larvae display aroused appetite for selected macronutrients in response to stimulation by food-related odors. Using this behavioral paradigm, we have identified a pair of higher-order neural circuits for perceptual processing of appetitive odor stimuli, each comprising a cluster of four DA neurons and a postsynaptic Dop1R1 neuron expressing NPF, an NPY-like neuropeptide (Figure 6). Our studies suggest that fly larvae offer a useful model for elucidating molecular and neural mechanisms underlying perceptual processing of appetitive olfactory and possibly other sensory cues.

### Contributions of DA signaling to larval perceptual judgment of appetitive odors

We have provided evidence that a subset of DA (DL2) neurons appears to make at least two major contributions to the perceptual processing of food-related olfactory stimuli. First, the DA neurons display excitatory responses to a broad spectrum of food-related odor stimuli, suggesting that they serve as a cognitive processing component for decoding and integration of chemically diverse food-related olfactory signals. Second, by genetic and pharmacological manipulations of the endogenous DA level, we have shown that the effective doses of odorants required for appetitive arousal must be significantly reduced in larvae pre-fed with L-dopa, or increased in larvae with reduced *TH* activity. These results suggest that the level of DA released from the DL2 neurons encodes both chemical identity and strength of an odor stimulus. Therefore, the DL2 neurons are essential for the compression of potentially multi-dimensional signals of food odors to one-dimensional DA codes.

### The Dop1R1 gating mechanism for shaping the inverted-U dose response

The dose-response analysis shows that the appetizing effects of both monomolecular and mixed odor stimuli follow an inverted U function. We have also provided evidence that the amount of DA released from the DL2 neurons is positively correlated with the strength of odor stimulation, and induction of appetitive arousal requires an optimum level of odor-evoked DA released from the DL2 neurons. For example, a higher non-appetizing dose of PA (e.g., 20μl PA) triggered a rapid and intense excitatory response from the DA neurons. As the result, a 30-sec stimulation by 20μl PA triggered appetitive arousal while its stimulation for 5 minutes failed to do so. Another key insight from this study is that odor-evoked DA signals encode no intrinsic appetitive values. Instead, larval perceptual recognition of appetitive odor cues requires a Dop1R gating mechanism, which is preset genetically, to selectively assign appetitive significance to a narrow spectrum of the DA signals. For example, a 50% reduction of Dop1R1 activity in *Dop1R1*/+ larvae led to a right-shift of the dose-response curve regardless whether the odor stimulus is monomolecular or chemically complex. Therefore, to be perceived as appetizing cues, odor-evoked DA signals must fall within a narrow range that matches the pre-existing level of Dop1R1 activity.

### The central role of NPF neurons in regulation of appetitive arousal

We have shown that two NPF neurons, located in the dorsomedial region of each brain lobe, are necessary and sufficient to elicit appetitive arousal in fed larvae. Anatomical and functional analyses also suggest that these NPF neurons likely form synaptic connections to the upstream clustered DL2 neurons ipsilaterally. In freely behaving fed larvae, the NPF neurons display excitatory responses to stimulation by appetizing doses of PA but not the non-appetizing doses, and such differential responses are gated by the Dop1R1 activity in the dorsomedial NPF neurons. Since dTRPA1-mediated genetic activation of the two dorsomedial NPF neurons, even under prolonged heat treatment, was sufficient to trigger increased feeding response, these results suggest that elevated NPF neuronal signaling plays an acutely role in the induction and maintenance of appetitive odor-aroused motivational state. It remains to be determined whether the NPF neurons are responsive to sugar stimulation. If so, this would provide evidence for an important role of these neurons in integrating DA-coded olfactory and sugar-evoked gustatory signals.

### Precision tuning of Dop1R1-gated NPF neuronal response to odor-evoked DA signals

We have provided molecular and cellular evidence for how the Dop1R1 activity in the NPF neurons shapes the inverted-U effects of appetitive odor stimuli. Using both behavioral and functional imaging assays, we show that Dop1R1 determines which range of odor-evoked DA signals may acquire appetitive significance through precision tuning of NPF neurons. Two separate genetic pathways have been identified for this process (Figure 6B). One of them, involving a Gβ13F/IRK2-mediated pathway, sets an upper limit of the optimum effective range of odor-induced DA signals by silencing the NPF neurons when Dop1R1is hyper-activated. Attenuation of the Gβ13F or IRK2 activity greatly broadens the effective dose range of DA signals, as evidenced by the change of the dose-response curve from an inverted U shape to a sigmoidal shape. Again, this result also strongly supports the notion that the ineffectiveness of stronger odor stimuli is caused by an excessive release of odor-evoked DA, which leads to a high level of dissociated G complex that activates IRK2 channels and subsequently silences the NPF neurons. The second Dop1R1/Gαs-mediated pathway provides a default excitatory mechanism that mediates NPF neuronal response to any odor stimuli that are at or above the minimal threshold strength. Therefore, they together define a two-layered precision tuning strategy.

The inverted-U effects of DA on cognitive functions have been widely observed in humans and other animals ^33,34^ Malfunctioned dopamine systems also underlie many psychiatric disorders such as schizophrenia ^35,36^. In the prefrontal cortex of mammals, an optimum level of D1-type DA receptor activity is crucial for spatial working memory, while its signaling at levels that are too low or too high leads to impaired working memory ^37–39^. Therefore, our findings raise the possibility that a homologous DA receptor-mediated tuning strategy may be used to mediate the inverted-U effects of DA in flies and mammals.

## Materials and Methods

### Fly Stocks and Larval Growth

All flies are in the w^1118^ background. Larvae were reared at 25°C, and early third instars (∼74 hr after egg laying, AEL) were fed before behavioral experiments as previously described^7^ (Wang et al., 2013). The transgenic flies include TH-GAL4 (Friggi-Grelin et al., 2003), UAS-*dTrpA1* (Hamada et al., 2008), UAS-*nsybGFP* ^19^, UAS-*sytGFP*^40^, UAS-*Denmark* (Nicolaï et al., 2010), UAS-GCaMP6.0(BL42749), UAS-*ple^RNAi^* (BL25796), UAS-*Gαs^RNAi^*(BL29576), UAS-Gβ76C^RNAi^(BL28507), UAS-Gβ13F^RNAi^(BL35041), UAS-*npf^RNAi^* (BL27237), UAS-*PKAi* (BL35550), UAS-*Arclight* (BL51056), UAS-*CaMPARI* (BL58761), UAS-*IRK2^RNAi^* (BL41981), UAS-*IRK1^RNAi^* (BL42644), UAS-*IRK3^RNAi^* (BL26720), and *npf-GAL4* were obtained from Bloomington stock center. UAS-Dop1R1^RNAi^ (V107058) was obtained from the Vienna Drosophila RNAi Center. *Dop1R1^f02676^* flies were described previously^15^. UAS-*CD4::spGFP^1-10^*, *LexAop-CD4::spGFP^11^* were kindly provided by K. Scott ^41^.

### Behavioral Experiments

Quantification of mouth hook contraction rate in liquid food was performed as previously described^20^. A published protocol for fly larvae odor stimulation was used with slight modifications^7^. Briefly, synchronized early third instars, fed on yeast paste, were stimulated for 5 minutes with specified doses of single or mixed odors including pentyl acetate (PA) (Sigma-Aldrich, 628-63-7), heptanal (Hep) (Sigma-Aldrich, 117-71-7), trans-2-Hexen-1-al (T2H) (Sigma-Aldrich, 6728-26-3), and the vapor of balsamic vinegar. After rinsing with water, larvae were tested for their feeding responses. Feeding media include agar paste (US Biological, A0940) containing 10% glucose, 0.5% Tryptone (Becton-Dickinson, 628200) or 3% oleic acid (Sigma-Aldrich, 112-80-1). UAS-*dTrpA1* was expressed by allowing larvae to feed in prewarmed yeast paste in a 31°C incubator for defined periods, followed by rinsing with 31°C water prior to feeding assays.

### 3IY and L-dopa Feeding

3-iodo-L-tyrosine (3IY) (Sigma-Aldrich, 70-78-0) and L-DOPA-ring-d3 (L-dopa) (Sigma-Aldrich, 53587-29-4) were used. The protocols for administration of 3IY and L-dopa were described previously^7^. The concentrations of 3IY and L-dopa in yeast paste were 10mg/ml and 0.5mg/ml, respectively.

### Molecular Cloning and Immunostaining

To construct the npf-LexA driver, a DNA fragment of ∼1-kb containing a region spanning from the 5’ regulatory sequence to the beginning of the second axon was amplified by genomic PCR. This fragment was subsequently cloned into the KpnI site in the pBPnlsLexA::GADflUw vector. Forward Primer: cagggagagagaacggagac; Reverse primer: gtgtcacaatgcaattgttcg. Tissue dissection and fixation, antibodies used and dilution conditions were described previously^7^.

### Targeted Laser Lesion

Protocols for calibration of 337 nm nitrogen laser unit and laser lesion experiments have been described^42^.

### CaMPARI Imaging

The conditions of larval feeding and odor treatment for CaMPARI imaging are identical to those for odor-aroused feeding behavioral assays. After odor stimulation, larvae were irradiated with PC light 405 nm LED array (200 mW/cm^2^, Loctite) for 5s. The treated larval CNS was dissected and individually scanned using a Zeiss LSM 510 confocal microscope.

### GCaMP imaging

The neural tissues of larvae were processed as previously described^7^. A Ca2+ indicator, GCaMP6s, was used for imaging odor responses by DA and NPF neurons. The odor delivery system involves a sealed 1.25L glass chamber fumigated with 7.5μl PA for 5 minutes. Odor was continuously delivered to larval head region by pumping at the rate of 0.28 L/min. The treated larval preparation was imaged using a Zeiss LSM 510 confocal microscope.

### Arclight imaging and data processing

The method for making larval CNS preparation is the same as previously described for calcium imaging preparation (7). The preparation was incubated in Drosophila PBS. Effective and ineffective odor vapors were prepared by fumigating a sealed 24L foam box with 150 or 800μl PA for 2 hours, respectively. Odor was continuously delivered to larval head region by pumping at the rate of 0.36L/min.

The protocol for ArcLight Imaging^25^ was followed with minor modifications. Briefly, larval CNS was imaged under 40X water immersion lens using Zeiss Axio Examiner. NeuroCCD-SM camera and Turbo-SM software (RedShirt Imaging) were used for recording and data processing. Images were captured at a frame rate of 100 Hz, and exposure time is 10ms. 2000 frames were collected for each of the seven 20s periods.

All the time series curves were low pass filtered with a Kaiser-Bessel 30 filter (200 Hz cut off). Then, each curve was fitted with a single exponential equation, I=Ae(-at). Bleaching of each curve was corrected based on the formula: It,corrected=It+(A----Ae(-at)). Normalization of each trace was achieved by dividing It,corrected with the average value. Standard deviation was obtained using normalized data.

### Statistical Analysis

Statistical analyses for behavioral and CaMPARI imaging assays were performed using One-way ANOVA followed by Dunnett’s or Sidak’s multiple comparisons test.

## SUPPLEMENTAL INFROMATION

Supplemental information includes Supplemental Experimental Procedures and four figures and one movie.

## AUTHOR CONTRIBUTIONS

Y.P. and M.M.M. performed behavioral assays. Y.P. and Y.Z. designed and performed imaging analyses. P.S., Y.P. and M.M.M. wrote the paper.

## ACKNOWLEDGMENTS

We thank J. Hirsh, K. Scott, F.W. Wolf, D.J. Anderson, the Bloomington Stock Center and Vienna Drosophila RNA I Center for fly stocks, and W. Neckameyer for anti-TH antibodies. We also thank P. Kner and K. Tehrani for help with processing of imaging data. This work is supported by grants from the U.S. National Institutes of Health (DK102209 and DC013333) to P.S.

## Supplemental Information

**Figure1--Figure supplement 1.**
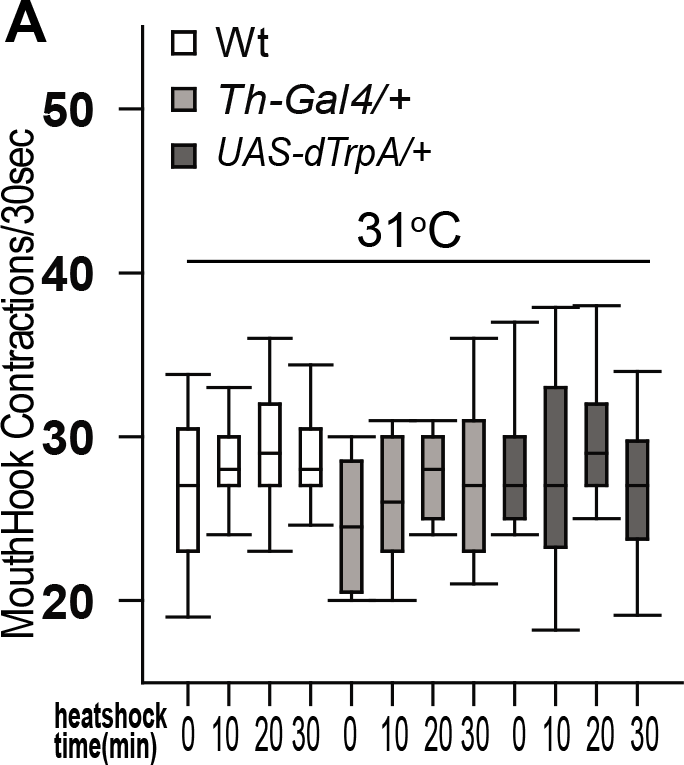
Control larvae display normal feeding responses after heat shock treatment. Fed larvae from three control groups display normal feeding responses after heat shocked at 31°C. n > 15.

**Figure3-- Figure supplement 2.**
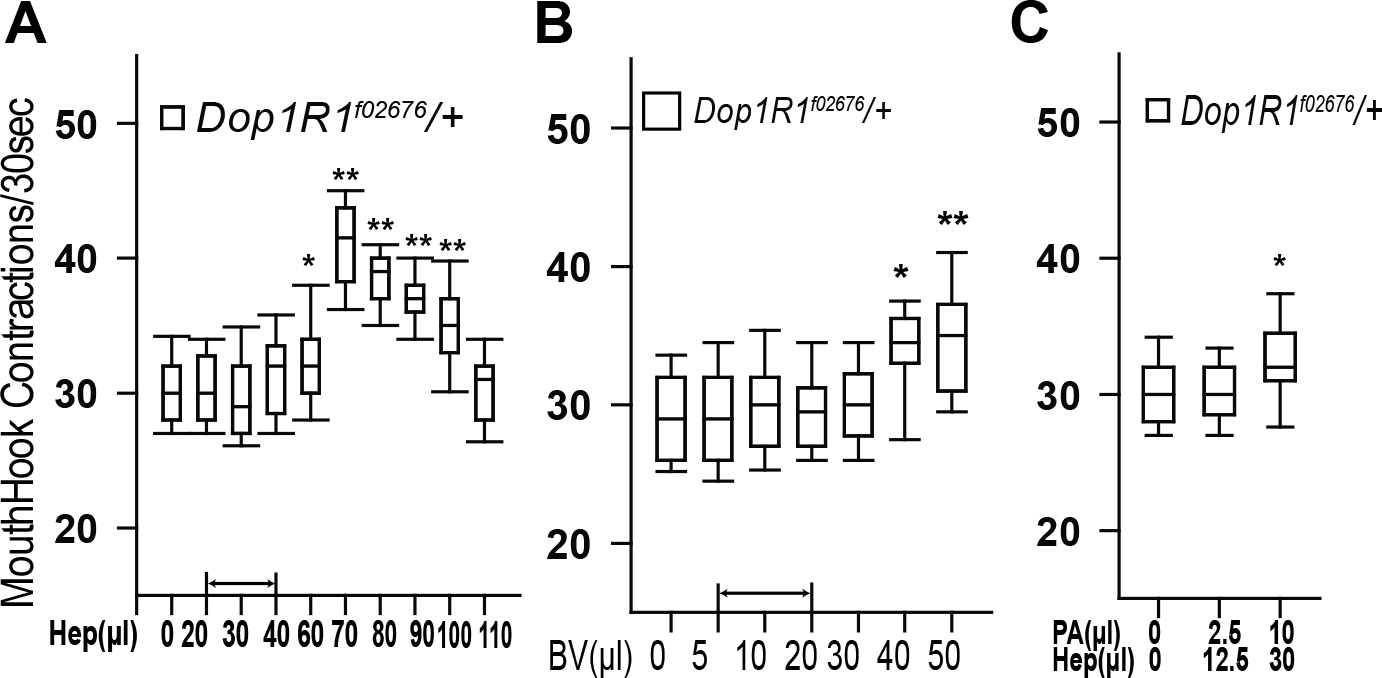
Heterozygous Dop1R1^f02676^ fed larvae showed right shifts in odor-aroused feeding responses (A, B) *Dop1Rl^f02676^*/+ fed larvae showed a right shift in the dose-response curve when stimulated by Hep or BV (n≥11). **(C)** *Dop1Rl^f02676^*/+ fed larvae failed to appetitive response to a normally appetitive binary mixture containing 2.5μl PA and 12.5μl Hep, but showed appetitive response to the normally non-appetitive binary mixture (n>16). *p<0.05;**P < 0.001.

**Figure4-- Figure supplement 3.**
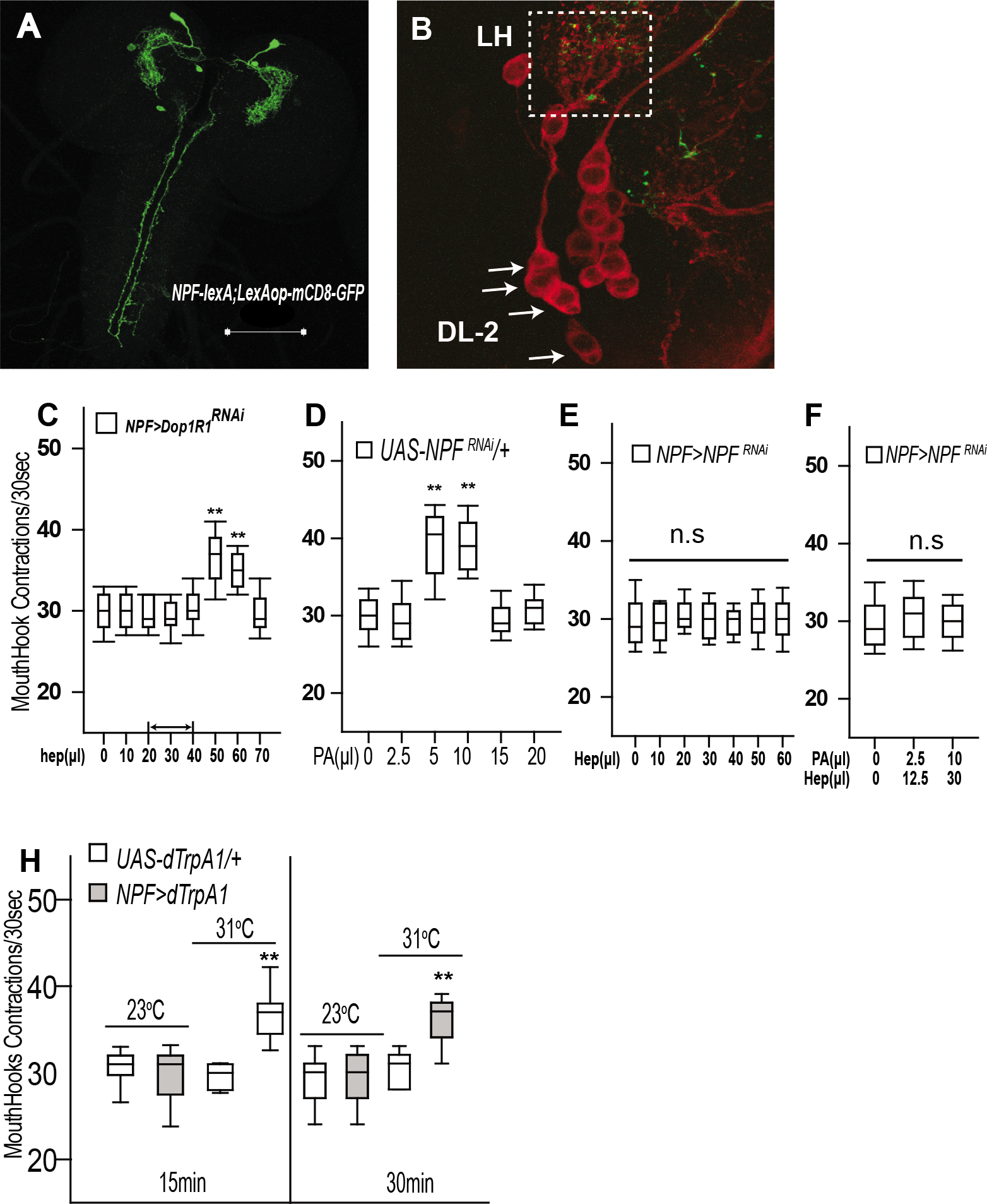
The odor-aroused feeding responses of Dop1R1- or NPF-deficient fed larvae. **(A)** The *NPF-LexA/LexAop-mCD8-GFP* larvae show the GFP expression predominantly in two dorsomedial NPF neurons. Scale bar=100μm. **(B)** A still image of the split GFP signals in the lateral horn region (from the first frame of Video 1). **(C)** *NPF-GAL4/UAS-Dop1R1^RNAi^* fed larvae displayed a right shift in the dose-response curve when stimulated with Hep (n>17). **(D)** UAS-*npf^RNAi^* fed larvae showed normal appetitive responses (n >14). **(E, F)** *NPF-GAL4/UAS-npf^RNAi^* fed larvae failed to show appetitive responses to all doses of Hep or binary mixtures tested (n>14). **(G)** Genetic activation of NPF neurons in NPF-GAL4/ UAS-*dTrpA1* fed larvae at 31°C for 15 or 30 min led to increased appetitive response (n>17). *P<0.05;**P < 0.001. **Figure4—Video 1 A 3D view of the presumptive synaptic connections between DA and NPF neurons**. The presumptive synaptic connections are labeled by split GFP (green) between DA neurons (red) and the NPF neuron in the lateral horn (DL2-LH, see Figure 4H).

**Figure5--Figure supplement 4.**
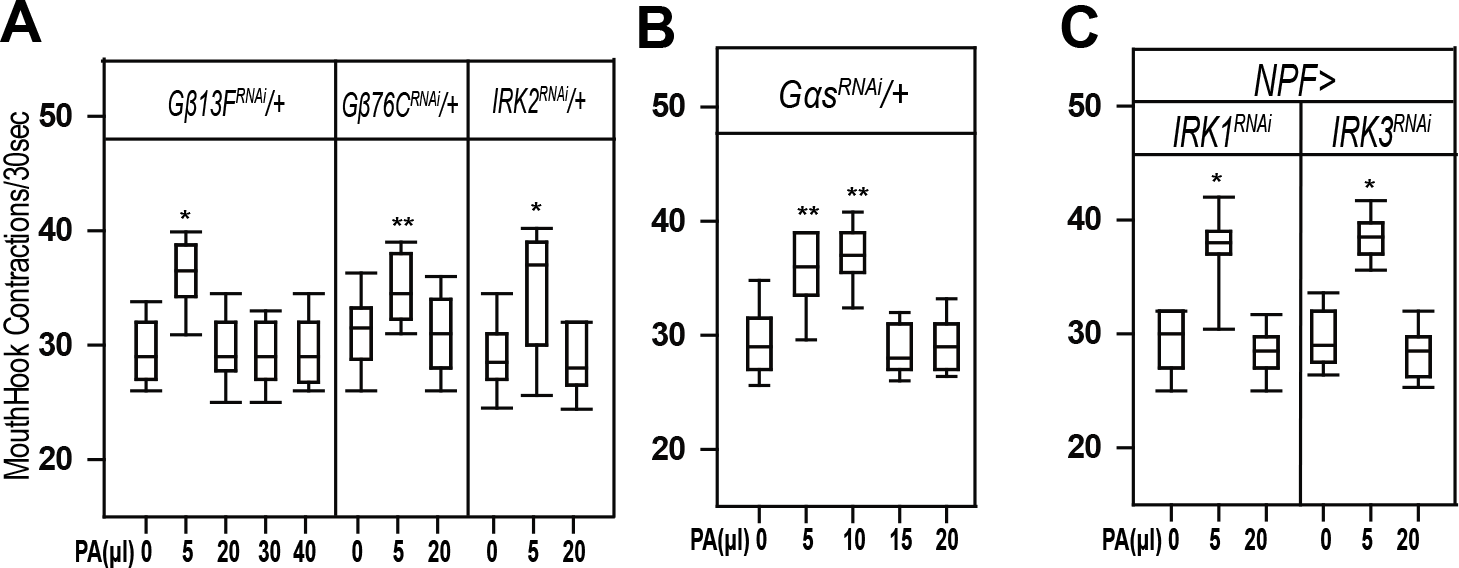
The effects of functional knockdown of Dop1R1 and its effectors on Odor-aroused Feeding Responses. **(A, B)** UAS-*Gβ13F^RNAi^*/+, UAS-*Gβ76C^RNAi^*/+ *and* UAS-*IR2^RNAi^*/+ control larvae showed normal feeding responses to PA. **(C)** PA-aroused UAS-*Gαs^RNAi^*/+ fed larvae displayed normal dose response. **(D)** NPF-GAL4/UAS-*IRK1^RNAi^* and NPF-GAL4/UAS-*IRK3^RNAi^* fed larvae displayed normal feeding responses to PA (n > 12). *p<0.05;**P < 0.001.

